# *Streptococcus equi* subsp. *zooepidemicus* associated with sudden death of swine in North America

**DOI:** 10.1101/812636

**Authors:** Matheus O. de Costa, Brad Lage

## Abstract

Historically described as a commensal of the swine upper-respiratory tract, *Streptococcus equi* subsp. *zooepidemicus* was only reported previously in Asia as an important swine pathogen. Here we report the isolation and whole genome characterization of *Streptococcus equi* subsp. *zooepidemicus* associated with a sudden death outbreak in pigs in North America.

## Text

*Streptococcus equi* subsp. *zooepidemicus* (*S. zooepidemicus*) is considered a commensal and opportunistic pathogen of several warm-blooded hosts, including humans, horses, different canines and swine. It is a Gram-positive, β-hemolytic coccus belonging to the Lancefield group C. It can cause severe disease characterized by pneumonia, septicemia and meningitis (1, 2). *S. zooepidemicus* has been suggested as a normal inhabitant of the palatine tonsils of pigs, being detected by both culture and high-throughput sequencing in samples collected from healthy animals (3). However, strains virulent to pigs have also been reported in the literature, particularly associated with high-mortality outbreaks of sudden death and respiratory disease in China (4). Currently, there are no vaccines available for this pathogen and control and prevention methods are hardly applied, given its commensal nature in swine. Here, we report an outbreak of sudden death associated with *S. zooepidemicus* in pigs housed in intensive rearing, commercial facilities in North America.

In April 2019 an outbreak of sudden death and abortions occurred in 4 loose-housed, commercial sow farms (approximately 9000 sows) in a large vertically integrated swine system within the province of Manitoba. This outbreak increased the cumulative mortality in the 3 affected sows herds by more than 1000 sows in the following 12 weeks. The abortion rate during this time period was approximately 11 x the normal rate. Animals were often described as apparently healthy during morning checks. Over the course of hours, sows would become unwilling to stand, develop fever, lethargy and die with no other apparent clinical signs. Other sows would abort and then go on to develop similar symptoms. Stressing factors in these farms, such as mixing of animals and the presence of other sick animals appeared to exacerbate outbreaks within pens. Animals were fed a commercial grade, nutritionally balanced diet as per ESF (electronic sow feeding) and had access to water *ad libitum*. Gross *post-mortem* examination of multiple animals, either euthanized or recently deceased, revealed the following common observations: rhinitis (mucopurulent discharge, mild, diffuse), pulmonary edema, gall bladder edema, hemorrhagic lymphadenopathy (tan to haemorrhagic) consisting of submandibular, cervical neck and bronchial lymph nodes, which taken together are suggestive of sepsis. All animals tested negative for PRRSV, *Mycoplasma hyopneumonia*e, SIV-A, PCV-3 and PCV-2 by real-time PCR. In parallel, Gram positive cocci were observed in imprints from heart and submandibular lymph nodes. Aerobic bacterial culture followed by Matrix-Assisted Laser Desorption/Ionization-Time Of Flight (MALDI-TOF) for identification of isolates revealed varying levels of *S. zooepidemicus* in liver, kidney, heart, brain, lung, spleen, and submandibular lymphnodes. Isolate identification was confirmed by two different veterinary diagnostic laboratories. Isolates (n=7, SAMN13058951, SAMN13058952, SAMN13058953, SAMN13058954, SAMN13058955, SAMN13058956, SAMN13058957) were found resistant to lincomycin, neomycin and tetracycline and susceptible to ampicillin, ceftiofur, penicillin and tilmicosin in a Kirby-Bauer disk diffusion assay.

DNA extraction was performed from isolates (DNeasy Powersoil Pro kit, Qiagen), quantified by Nanodrop (3300) and PicoGreen (Quant-iT dsDNA) and processed for sequencing (Illumina Nextera XT library prep kit). Sequencing was performed using MiSeq Nano V2 (2×250 paired-end). Samples yield an average of 149,017 high quality reads, suggesting 50x coverage (genome size averaged 2.1 mbp). Genome assembly, annotation and downstream analyses were conducted using the PATRIC package (5). Genomes averaged 2.1 million bp in size, and 41.34% in GC content. All isolates were similar to previously published *S. zooepidemicus* genomes (Figure 1), demonstrating a whole-genome average nucleotide identity (ANIscore) of 99.7% to strain *S. zooepidemicus* ATCC35246. This particular strain was reported as isolated from a septicaemic pig during an outbreak that killed over 300,000 pigs in Sichuan province, China, in 1976 (6). Interestingly, all isolates had an average ANIscore of 97.3%, when compared to *S. zooepidemicus* strain 4047, an isolate considered virulent, obtained from a horse diagnosed with strangles in the United Kingdom (7). In addition, all isolates obtained from pigs, regardless of what outbreak, were profiled as MLST (multi-locus sequence type) ST-194, including strain ATCC35246. Antimicrobial resistance genes identified in isolates from this outbreak included *gidB, S12p* (streptomycin), *rpoB* (rifampin), *S10p* (tetracycline), *kasA* (triclosan), *PgsA, LiaR, LiaS* (daptomycin), *folA, Dft* (trimethoprim), *folP* (sulfadiazine) and *FabK* (triclosan). Virulence factors found included the previously described *szm, lmb, fbpZ, skc, has* operon and *mag* regulon, which help explain the highly-virulent lifestyle of these isolates.

**Figure 1.**
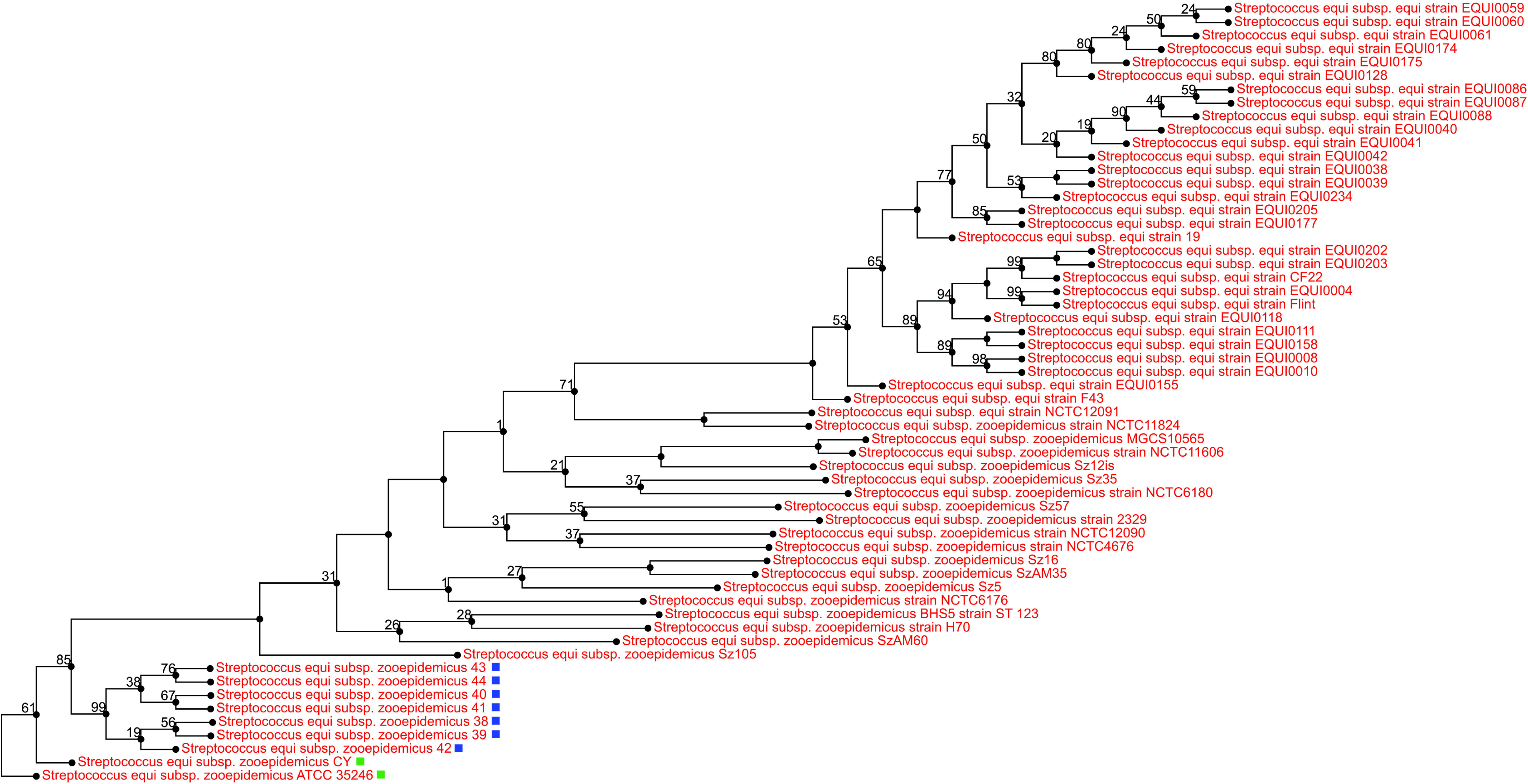
Phylogenetic tree (all-shared proteins) of *Streptococcus equi* subsp. *zooepidemicus* whole-genome sequences obtained from the reported outbreak in pigs from North America (blue blocks, PRJNA578379), compared with previously published human, dog, horse and pig (green blocks) sequences from GenBank (n=58). Tree inferred using BLAST followed by FastTree within the PATRIC package(5). Support values shown indicate the number of times a particular branch was observed in the support trees using gene-wise jackknifing.

Taken together, these findings suggest the emergence of *S. zooepidemicus* ST-194 as a cause of mortality in pigs in North America. This specific sequence type seem to be particularly virulent to pigs, for reasons that remain unexplained. Given the clinical presentation described here, this pathogen requires special attention and should no longer be overlooked, due to its historically accepted commensal lifestyle, when conducting diagnostic investigations.

## Acknowledgments

The authors would like to acknowledge the Manitoba Veterinary Diagnostic Services for assistance.

## Author Bio

Dr. Costa is an Assistant Professor at the University of Minnesota, USA. His research interests include swine bacterial pathogens to which no preventive or control methods are available besides antibiotics. Dr. Lage is the head Veterinarian at Maple Leaf Agri-Farms, the swine division for Maple Leaf Foods based in Landmark, Manitoba.

